# Jagged-1+ Skin Tregs Modulate the Innate Immune Response to Wound Healing

**DOI:** 10.1101/2024.06.10.598375

**Authors:** Prudence PokWai Lui, Jessie Z. Xu, Hafsah Aziz, Monica Sen, Niwa Ali

## Abstract

Skin-resident regulatory T cells (Tregs) play an irreplaceable role in orchestrating cutaneous immune homeostasis and repair, including the promotion of hair regeneration via the Notch signaling ligand Jagged-1 (Jag1). While skin Tregs are indispensable for facilitating tissue repair post-wounding, it remains unknown if Jag1-expressing skin Tregs impact wound healing. Using a tamoxifen inducible Foxp3^creERT2^Jag1^fl/fl^ model, we show that loss of functional Jag1 in Tregs significantly delays the rate of full-thickness wound closure. Unlike in hair regeneration, skin Tregs do not utilize Jag1 to impact epithelial stem cells during wound healing. Instead, mice with Treg-specific Jag1 ablation exhibit a significant reduction in Ly6G+ neutrophil accumulation at the wound site. However, during both homeostasis and wound healing, the loss of Jag1 in Tregs does not impact the overall abundance or activation profile of immune cell targets in the skin, such as CD4+ and CD8+ T cells, or pro-inflammatory macrophages. This collectively suggests that skin Tregs may utilize Jag1-Notch signalling to co-ordinate innate cell recruitment under conditions of injury but not homeostasis. Overall, our study demonstrates the importance of Jag1 expression in Tregs to facilitate adequate wound repair in the skin.

## INTRODUCTION

The skin, the largest mammalian organ, acts as a critical physical barrier from constant environmental traumas. A diverse array of immune and non-immune cell types reside in or migrate to the skin during pre- and post-natal development, influenced by the local inflammatory microenvironment. Over the past decade, skin resident regulatory T cells (Tregs), a major T cell population, have been implicated in various processes, including hair regeneration^1,2^, full-thickness wound healing^3^, epidermal barrier repair^4,5^ and fibrosis^6^. These functions are facilitated by the immunosuppressive capacity of skin Tregs, including restriction of pro-inflammatory cytokine production and myeloid cell accumulation^3,4^, as well as their interaction with non-immune tissue cells such as epithelial keratinocytes^5^, fibroblasts^6^ and stem cells^1,4^.

Multiple single cell studies have illustrated tissue Tregs are transcriptionally distinct from those within secondary lymphoid organs^7–10^. The site where Tregs are seeded determines their phenotype and shapes their functions^7,10^. Previous RNA sequencing analysis identified Jagged-1 (Jag1), one of the five Notch signalling ligands, as a key transcript preferentially expressed in skin Tregs compared to skin-draining lymph node (SDLN) Tregs^1^. Interestingly, intraperitoneal injection of IL-2/anti-IL2 antibody complexes can selectively expand Jag1+Tregs in murine skin but not in SDLNs^11^. *In silico* data indicate that skin Tregs are also phenotypically distinct from other tissue-resident Tregs^6,9,10^. Ligand-receptor prediction analyses from scRNA-seq datasets further suggest that skin Tregs likely interact with epithelial cells and hair follicle stem cells via Jag1-induced Notch signalling^2,5^, underscoring the potential importance of Jag1 in skin Tregs. Yet, the question of whether Jag1 expression is indeed unique to skin Tregs remains unanswered.

Notch signalling is crucial for hair follicle differentiation and postnatal maintenance homeostatically^12^, as well as in regulating wound healing^13^. Conditional deletion of Jag1 in epidermal stem cells (K15^cre^Jag1^fl/fl^) delays wound closure, whereas the addition of Jag1 peptide accelerates wound healing^13^, demonstrating the significance of Jag1 in the wound repair process. Although it has been established that Jag1 expressed on non-lymphoid cells can drive T cell fate decisions^14,15^, promote Treg expansion^16–18^, and that Notch signalling regulates Treg immunosuppressive capacity^19^, the function of Treg-derived Jag1 remains largely unknown. Previously, we have shown the perturbation of Jag1 in skin Tregs hinders hair follicle stem cell (HFSC) proliferation, delays the induction of the hair growth phase, and ultimately impedes hair regeneration^1^. Whether the relationship between Jag1+ Tregs and hair regeneration translates to other skin Treg-mediated mechanisms, such as wound healing, remains unexplored.

Here, we report that Jag1 is preferentially expressed in skin Tregs compared to Tregs residing in other tissues. Despite Jag1+Tregs displaying higher CTLA4 and CD25 surface expression, the absence of Jag1 in Tregs does not alter skin integrity nor overall cutaneous immune dynamics during homeostasis. However, during full thickness wound healing, mice with Jag1 deficiency in Tregs heal significantly slower than controls. Unlike in hair regeneration, skin Tregs in wounded mice do not utilize Jag1 to alter HFSC activation but rather promote neutrophil accumulation at the wound site. This study sheds light on an alternative function of Jag1 in skin Tregs in facilitating adequate cutaneous wound repair.

## RESULTS

### Jagged1 is a Skin Treg Preferential Marker

To determine whether Jag1 is uniquely expressed in skin Tregs, we began by re-analysing a bulkRNAseq dataset focusing on Tregs from various tissues^10^. Our analysis confirmed that skin Tregs exhibited the highest levels of *Jag1* compared to Tregs from blood, spleen, and other organ tissues (visceral adipose tissue (VAT), lung and colon) (Figure 1A). Differential expression analysis using edgeR and limma packages normalized counts and compared the adjusted p-values (Fig1B) and log fold changes (Fig1C) of Jag1 expression in skin Tregs against other tissue Tregs. We included Cd45 (Ptprc) and common Treg markers (Foxp3, Ctla4, Cd25, and Icos), along with known tissue Treg-associated transcripts (Gata3 and Areg) as controls.

**Figure 1.**
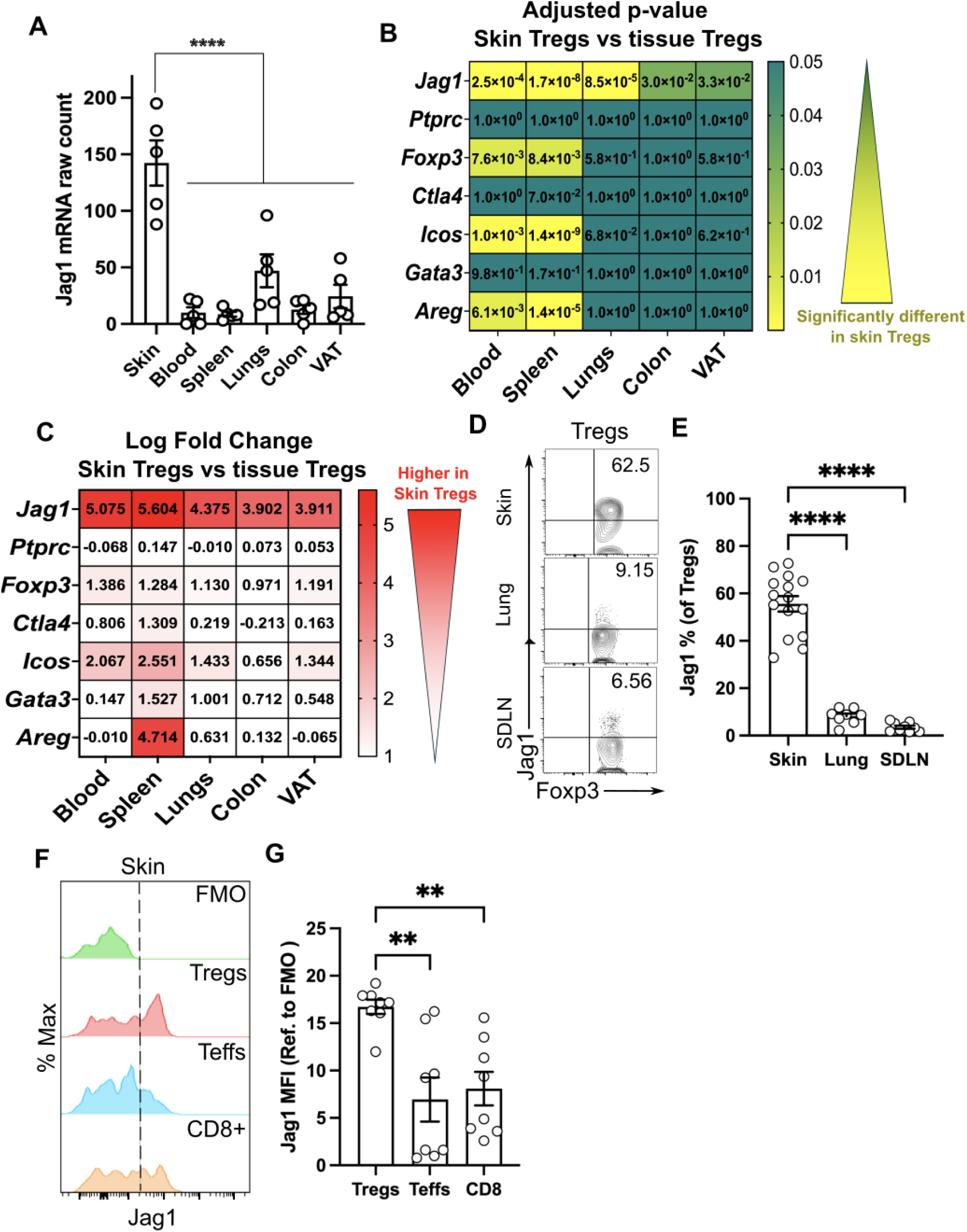
Jagged1 is preferentially expressed in skin Tregs. **(A)** Jag1 mRNA counts from bulk RNAseq of Tregs from different tissue sites. (n = 5). (**B** and **C**) Heatmap showing the adjusted p-value (B) and log fold change (C) of genes expressed in Tregs from skin compared against those from other tissues. (**D** and **E**) Representative flow cytometry plot and quantification (E) of Jag1+ Tregs in skin, lung and skin-draining lymph nodes (SDLNs) (n = 8-15). (**F** and **G**) Representative histogram (F) and quantification (G) of Jag1 expression in different skin T-cell populations (n= 8). Data in (D,E) or (F,G) were pooled from 3 independent experiments. Individual data points are shown and presented as mean ± SEM. Statistics in (A), (E) and (G) were calculated by one-way ANOVA, ** *p <*0.01, **** *p* < 0.0001.

Notably, *Jag1* expression in skin Tregs was significantly and consistently higher than, not only those in blood or lymphoid organs, but also Tregs residing in lung, colon and VAT (Figure 1B&C). Skin Tregs expressed an average 4.5-fold higher level of *Jag1* than other surveyed Tregs, clearly distinguishing Jag1 from other Treg-associated genes. As expected, *Cd45 (Ptprc)* showed no significant differences (high adjusted p-value and low log fold change) between skin and other tissue Tregs. Skin Tregs expressed similar level of *Foxp3* and common Treg markers, as well as tissue Treg-associated genes, compared to Tregs in the lung, colon, and VAT. While *Ctla4* and *Gata3* expression levels remained similar, *Foxp3* and *Icos* were significantly but mildly upregulated in skin Tregs, when compared to those in blood or spleen. Similar to *Jag1*, *Areg* was highly expressed in skin Tregs compared to splenic Tregs, but not when compared to other tissue Tregs. Collectively, this highlights Jag1 as a differentially expressed marker in skin Tregs.

To further validate these findings, we performed flow cytometry on Tregs from skin, lung and SDLN. Around 55.59% (±12.61%) of skin Tregs expressed Jag1, in contrast to only 8.18% (±3.20%, p < 0.0001) in lung Tregs and 3.54% (±1.90%, p < 0.0001) in SDLN Tregs (Figure 1D & E, gating strategy in Supplementary Fig1A). The mean fluorescence intensity of Jag1 was significantly higher in skin Tregs (16.72 ± 2.15), in comparison to other CD4+Foxp3- T effector (Teffs) (6.95 ± 6.57, p = 0.0014) and CD8+ T cells (8.10 ± 5.01, p = 0.0041), indicating preferential expression of Jag1 in skin Tregs.

We then examined whether these abundant Jag1^pos^ skin Tregs are phenotypically distinct from non-Jag1-expressing (Jag1^neg^) skin Tregs. In adult mice, Jag1^pos^Tregs showed significantly higher proportions and levels of CTLA4 (Supplementary Fig2A-D) and CD25 (Supplementary Fig2E-H), but not ICOS (Supplementary Fig S2I-L). Jag1^pos^ skin Tregs co-expressed CTLA4 and CD25 more frequently than Jag1^neg^Tregs (Supplementary Fig2M&N, 37±4.64% vs 23.5±7.90%, p = 0.0312), suggesting that Jag1^pos^Tregs may be phenotypically more active.

### Jag1^pos^ Tregs are Dispensable During the Steady State

Under homeostatic conditions, skin Tregs suppress long-term CD8 and Teff cell-driven hair follicle-associated inflammation via CD25^Ref.20^. Given that Jag1^pos^ skin Tregs express higher levels of CD25 than Jag1^neg^ skin Tregs, we hypothesised that Jag1^pos^ skin Tregs may be functionally important during the steady state. To test this, and to avoid influencing early skin development, we created a tamoxifen inducible cell specific model to delete Jag1 in Tregs. This involved crossing mice expressing EGFP-creERT2 gene under the Foxp3 promoter^21^ to mice carrying a Jag1 floxed allele with a dysfunctional Delta-Serrate-Lag2 domain of Jag1^22^ to generate Foxp3^creERT2^Jag1^fl/fl^ and Foxp3^creERT2^Jag1^fl/wt^, hereafter denoted as “Foxp3^ΔJag^^1^“ and “Foxp3^Ctrl^”. Following intraperitoneal injection of tamoxifen (Figure 2A), we observed effective downregulation of Jag1 transcript in sorted Tregs from Foxp3^ΔJag1^ compared to Foxp3^Ctrl^ mice (Supplementary Figure3A). Expression of *Jag1* in Teffs, CD8+ T cells, and *Foxp3* expression in Tregs (Supplementary Figure3B) showed no differences between Foxp3^ΔJag^^1^ and Foxp3^Ctrl^ animals, indicating Treg-specificity of Jag1 deletion in this model.

**Figure 2.**
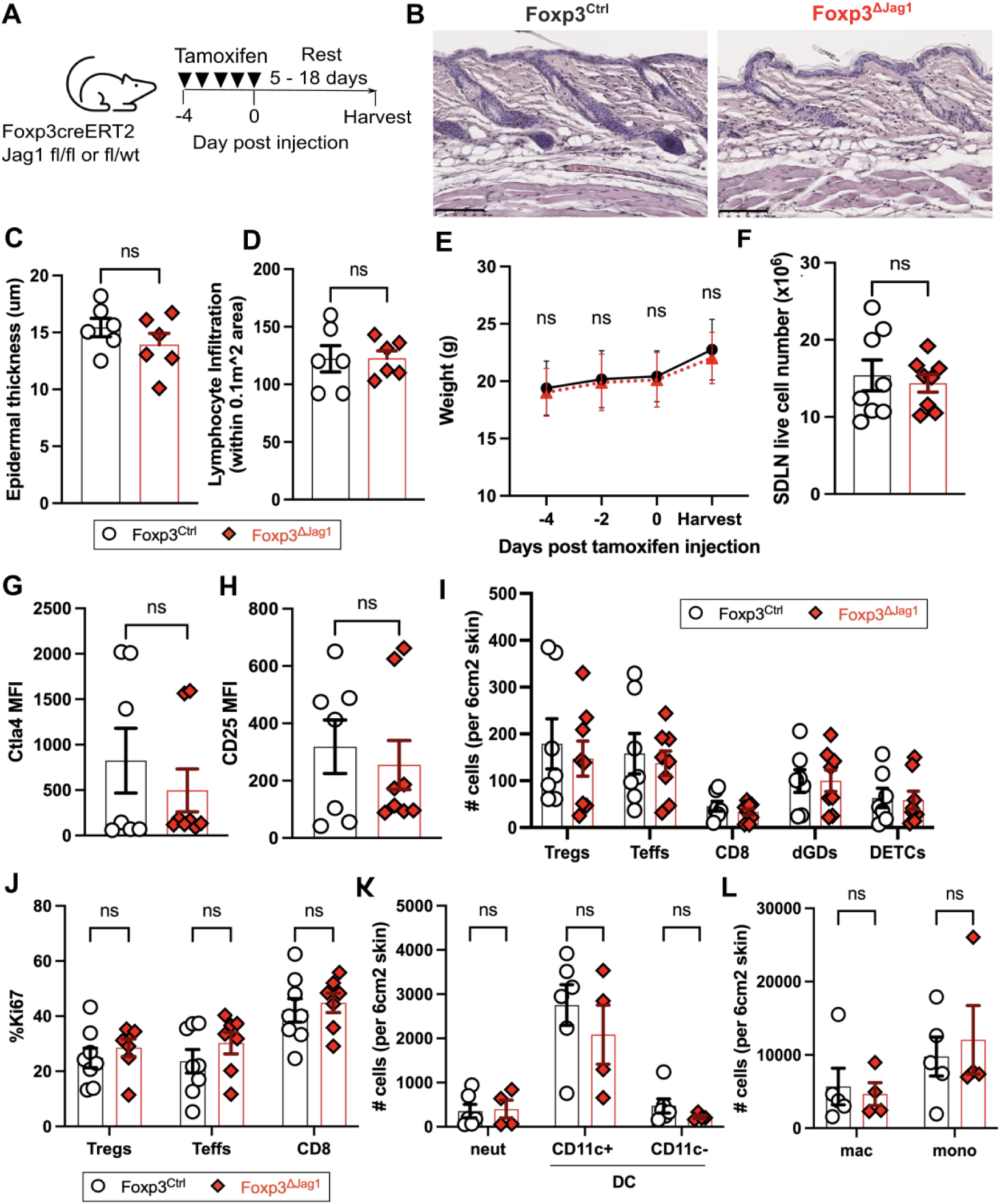
Jag1 in Tregs is dispensable for skin homeostasis. (**A**) Experimental schematic illustrating intraperitoneal injection of 100mg/kg dose of tamoxifen daily for 5 days, with 5-18 days rests before harvest for downstream analysis. (**B**) Representative H&E staining of Foxp3^Ctrl^ and Foxp3^ΔJag1^ skin. Scale bars represent 100μm. (C and D) Quantification of epidermal thickness (C) and lymphocyte infiltration (D) from three regions-of-interest in H&E staining of Foxp3^Ctrl^ and Foxp3^ΔJag1^ skin (n = 2 per group)(E) Average weight of treated animals traced throughout the course of tamoxifen injection and harvest end point (n = 4 per group). (F) Total live cells from Foxp3^Ctrl^ and Foxp3^ΔJag1^ SDLNs (n = 8 per group). (G and H) Flow cytometric quantification of mean fluorescence intensity (MFI) of CTLA4 (G) and CD25 (H) in Tregs from Foxp3^Ctrl^ and Foxp3^ΔJag1^ skin (n = 6-7). (I and J) Quantification of skin T cell abundance (I) and proliferation (J) in Foxp3^Ctrl^ and Foxp3^ΔJag1^ mice (n = 6-7 per group). (K and L) Quantification of neutrophils (neut), CD11c+ and CD11c-dendritic cells (DCs), macrophages (mac) and monocytes (mono) from wildtype Foxp3^Ctrl^ and Foxp3^ΔJag1^ mice (n = 4-6 per group). Data in (E) to (L) were harvested from 4 independent experiments collected between 5 - 18 days post last tamoxifen injection. Data in (B to D) were from 1 and (K and L) from 2 of these 4 experiments. Each individual data point represent one biological replicate. Results were presented as mean ± SEM. Statistics were calculated by unpaired t-test (C, D, F, G and H) and two-way ANOVA (I, J, K and L), ns = non-significant.

Despite significant systemic loss of Jag1 mRNA in Tregs, the absence of Jag1 did not trigger skin inflammation. H&E staining revealed similar morphology (Figure 2B), epidermal thickness (Figure 2C) and lymphocytic infiltration (Figure 2D) in Foxp3^ΔJag^^1^ and Foxp3^Ctrl^ animals. Body weight throughout tamoxifen injections (Figure 2E) and SDLN live cell numbers (Figure 2F) remained unchanged between animals with or without Jag1 in Tregs during the steady state. Loss of Jag1 in Tregs did not alter overall CD25 or CTLA4 expression in skin Tregs (Figure 2G&H), nor the abundance and proliferation of skin-resident Tregs, Teff, CD8, gamma-delta T cells (Figure 2I &J) and other myeloid populations (Figure 2K&L, gating strategy in Supplementary Fig1B). Together, this suggests that Jag1 expression in Tregs is dispensable for maintaining skin immune homeostasis.

### Jag1^pos^ Tregs are Highly Activated During Early Wound Healing

One of the most common skin traumas is cutaneous injury. Treg depletion hinders both epithelial restoration^4^ and the full-thickness wound healing process^3^. One known function of Jag1^pos^Tregs is their role in facilitating hair follicle stem cell (HFSC) proliferation during hair growth phase transitions^1^. Therefore, we investigated whether Jag1^pos^ Tregs are involved in wound healing. Wound healing comprises four major overlapping stages: haemostasis, inflammation, proliferation, and remodelling^23^. Skin Treg abundance at full-thickness wound sites peaks at 7 days post wounding (dpw) and returns to homeostatic levels by 14dpw^3^. In Foxp3-DTR mice that permits systemic loss of all Tregs, delayed wound healing is observed only when Tregs are depleted during the inflammation phase, but not later. Jag1 expression in whole skin lysates follows a similar dynamic, being upregulated during the first 7dpw and downregulated from 14dpw^24,25^. Thus, we hypothesised that the first 7 days are likely to be a crucial period in which Jag1^Pos^ Tregs may have an impact.

We first characterised the dynamic expression of Jag1+Tregs by creating two 4mm full-thickness wounds on the dorsum of wildtype adult mice at 0 days post wounding (dpw) and harvesting skin around the wound at inflammatory (2dpw), proliferative (5dpw) and remodelling (12dpw) phases (Figure 3A). Notably, 25%-48% of skin Tregs expressed Jag1 during the first 5 days post-wounding, in contrast to 6.35% (±2.25%, p = 0.0014) at 12dpw (Figure 3B&C). At 5dpw, Jag1^pos^+Tregs were significantly more proliferative (Figure 3D&E) and co-expressed higher levels of both CTLA4 and CD25 activation markers (Figure 3F&G), relative to Jag1^neg^Tregs. In contrast, no differences in Ki67 or CTLA4+CD25+ proportions were observed between Jag1^pos^ and Jag1^neg^ skin Tregs at 2dpw, indicating that 5dpw is a critical period for Jag1+ Tregs’ functional prominence.

**Figure 3.**
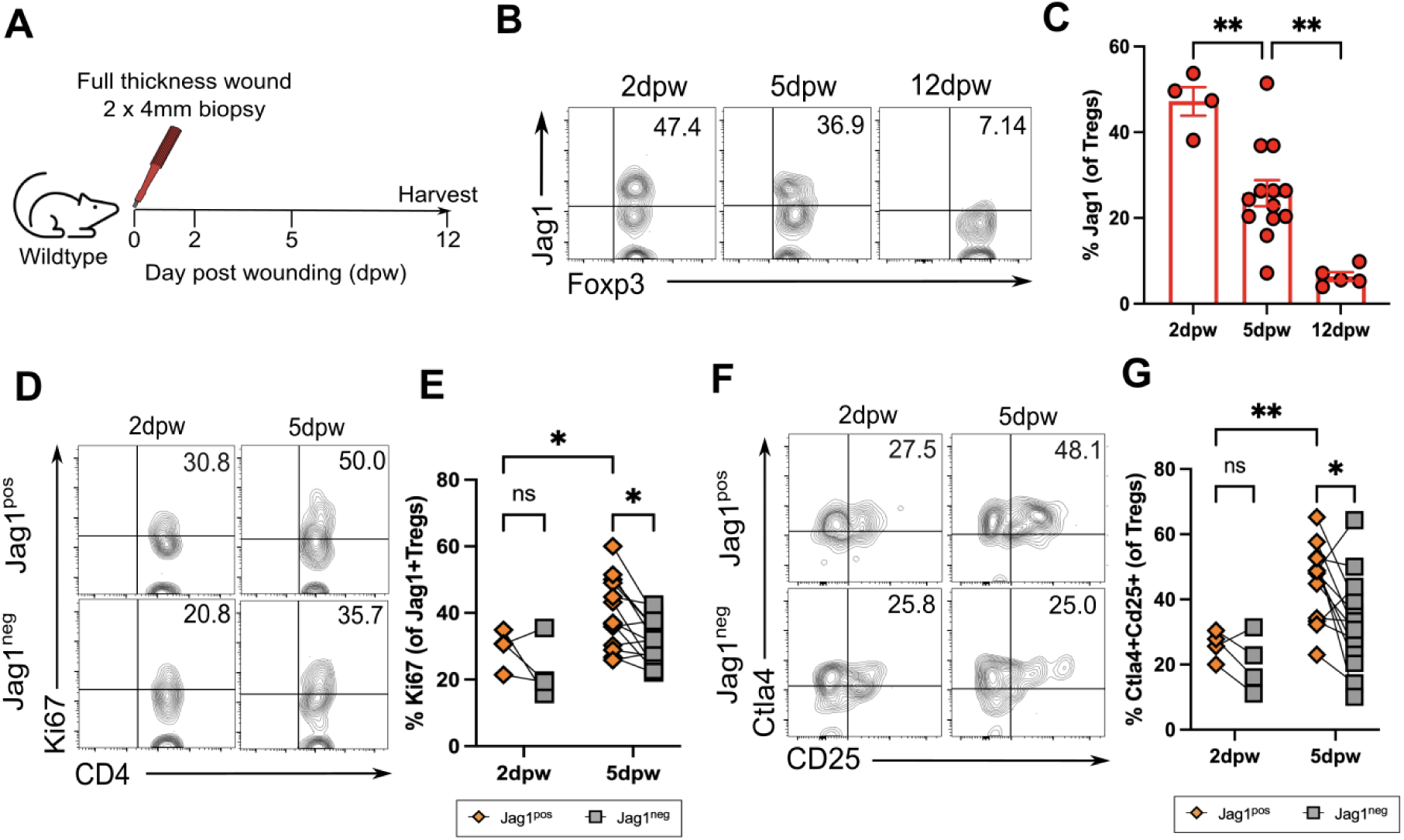
Jag1+Tregs are most abundant and activated at 5 days post wounding. **(A)** Experimental schematic in which two circular full thickness wounds were generated on the dorsum of wildtype mice using a 4mm punch biopsy. Skin was harvested at 2 days post wound (dpw), 5dpw and 12 dpw. (**B** and **C**) Representative flow plots and quantification of % Tregs expressing Jag1 in wounded skin at 2dpw, 5dpw and 12dpw (n = 4-12 per group). Representative flow plots and quantification of (**D** and **E**) %Ki67 and (**F** and **G**) %CTLA4+CD25+ in Jag1+ Tregs and Jag1-Tregs from 2dpw and 5dpw wounded skin (n = 4-15 per group). Data in (B) to (G) were pooled from 2 independent experiments. Results were presented as individual data points with mean ± SEM in (C) and paired data-points in (E and G). Statistics were calculated by one-way ANOVA (A) and two-way ANOVA (E and G), * *p <* 0.05, ** *p <* 0.01, ns = non-significant.

### Jag1+ Tregs Promote Wound Healing

We then assessed the role of Jag1^pos^Tregs in wound healing by creating two full thickness excisional wounds of the dorsal skin of Foxp3^ΔJag^^1^ and Foxp3^Ctrl^ mice, and measuring the wound healing rate over time (Figure 4A). Mice lacking Jag1-expressing Tregs showed delayed wound closure compared to wildtype Foxp3^Ctrl^ mice treated with tamoxifen (Figure 4B & C). The most pronounced difference was observed at 5dpw, with 78.4% (± 9.3%) wound closure in wildtype Foxp3^Ctrl^ mice versus 66.2% (±15.9%) in Foxp3^ΔJag^^1^ animals (Figure 4D, p = 0.0041). Histologically, Foxp3^ΔJag^^1^ skin showed a trend towards thicker epithelial wound edges (Figure 4E, red arrow), more keratinocyte precipitation at wound edge (Figure 4E,black arrow) and disorganisation between epithelial layer and pus cells (Fig 4E, white arrow). These results illustrate skin Tregs require Jag1 for effective wound closure particularly during the early (5 dpw) proliferation phase.

**Figure 4.**
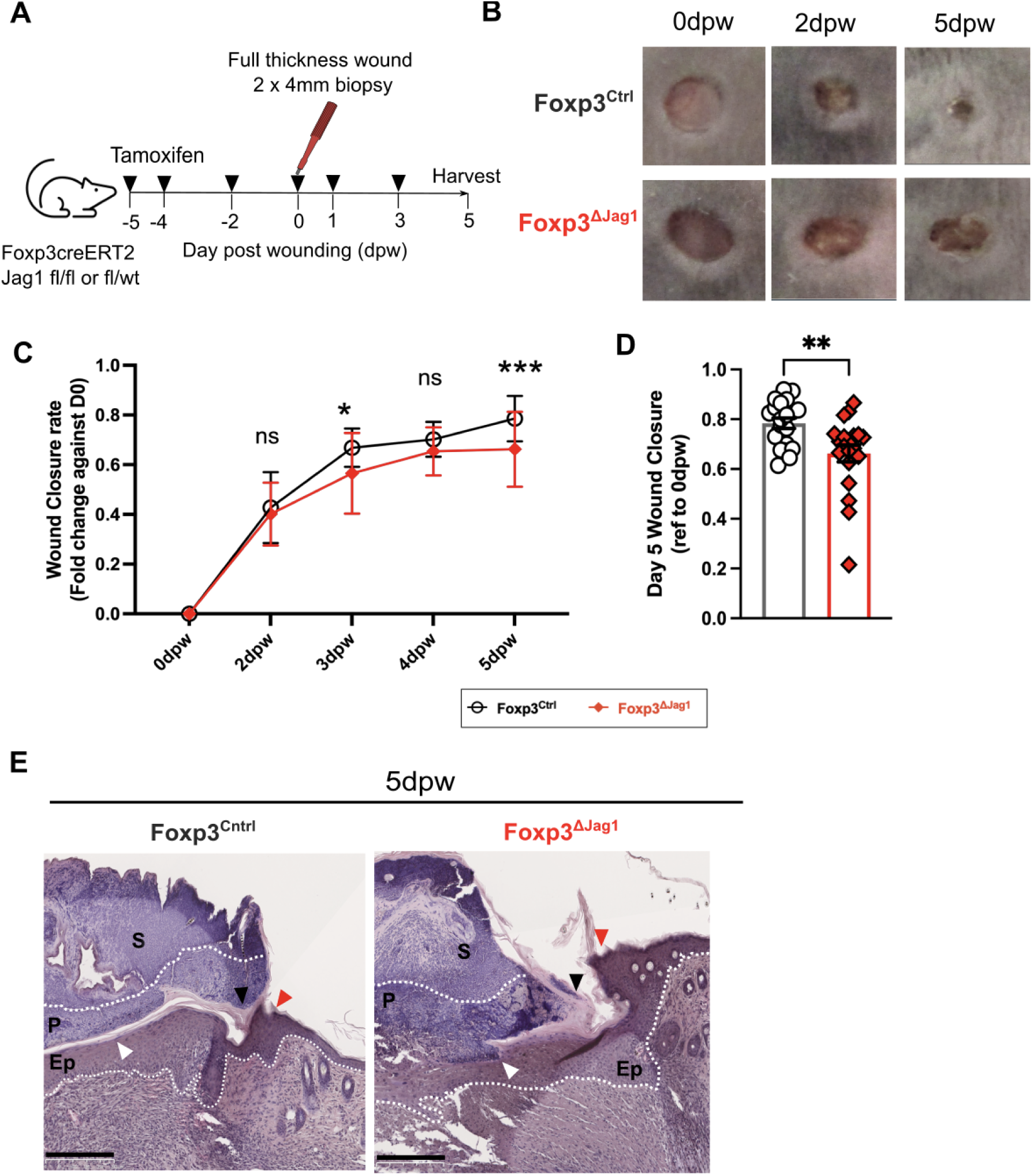
Jag1+ Tregs are required for wound healing. **(A)** Experimental schematic in which Foxp3^ΔJag1^ and “Foxp3^Ctrl^ mice were injected with 4 doses of tamoxifen, before wounding with 4mm biopsy punches. Two more doses of tamoxifen were given thereafter before wounded skin were harvested at 5dpw. **(B)** Representative clinical images of the healing progression at 0dpw (immediate after wounding), 2dpw, and 5dpw **(C** and **D)** Quantification of wound closure, calculated by fold change against wound area at 0dpw, with (C) showing the kinetics throughout the experiment, and (D) ratio at 5dpw (n = 20-22 per group). (E) Representative H&E image of Foxp3^Ctrl^ and Foxp3^ΔJag1^ wounded skin harvested at 5dpw, with labels of scab (S), pus cells (P) and epidermis (Ep). Arrows illustrated region of interest, including epidermal wound edge (red), keratinocyte precipitation (black) and border between pus cells and epidermis under scab (white). Scale bars represent 250μm. (n = 4 per group). Data were pooled from 4 independent experiments. Results were presented as mean ± SEM in (C) and individual biological replicates as each data point with mean ± SEM in (D). Statistics were calculated by two-way ANOVA in (C), and unpaired t-test in (D).* *p <* 0.05, **p < 0.01, *** p < 0.001.

### Jag1+ Tregs Modulate Neutrophil Accumulation During Early Wound Healing

We next explored the mechanisms by which skin Tregs use Jag1 to orchestrate wound closure. Upon tissue challenge, skin Tregs primarily perturb inflammation through two mechanisms: either via conventional immunosuppressive capacity^3,4^ or enhance tissue repair via cross-talk with epithelial cells and HFSCs^1,4^. Dysregulation of skin Tregs leads to unwanted inflammation, mainly contributed by excessive accumulation and proliferation of neutrophils, pro-inflammatory macrophages, CD4+, and CD8+ T cells during homeostasis ^20^, epidermal injury^4^, and full thickness injury^3^. Treg-specific loss of Rbpj, a downstream transcription factor of canonical Notch signalling, has been shown to mediate upregulation of Foxp3, CTLA4 and CD25 expression in splenic Tregs during homeostasis^19^. With Jag1^pos^Tregs co-expressing higher level of CTLA4 and CD25 during both homeostasis and wound healing, relative to Jag1^neg^ Tregs, we questioned whether the absence of Jag1 in Tregs can affect the overall immunosuppressive ability of skin Tregs. We reasoned this may impact the accumulation of T and/or myeloid cells during wound healing and could influence wound closure through regulating local inflammation.

Similar to the steady state (Figure 2), the absence of Jag1 in Tregs did not affect overall Treg abundance, as quantified by both flow cytometry and immunofluorescence staining (Supplementary Figure 4A&B). Proliferation and Foxp3 expression in skin Tregs also remained unaffected (Supplementary Figure 4C&D). None of the Treg phenotypic attributes were altered by loss of Jag1, including the proportion of skin Tregs co-expressing CTLA4+CD25+ (Supplementary Figure 4E), CD25 (Supplementary Figure4F) or ICOS (Supplementary Figure4G). IL33 is an alarmin highly induced in keratinocytes in response to cutaneous wounding^26^. IL33 receptor (ST2) is highly expressed in skin Tregs^10^, and recently found crucial for suppressing bleomycin-induced skin fibrosis^27^. However, we found that Tregs from Foxp3^ΔJag^^1^ skin did not alter ST2 expression level either (Supplementary Figure4H). Intriguingly, Jag1 ablation in Tregs resulted in a mild but significant reduction in CTLA4 levels (Supplementary Fig4I). Yet, both the accumulation and proliferation of Teffs and CD8+ T cells remained unchanged between Foxp3^ΔJag^^1^ and Foxp3^Ctrl^ skin (Supplementary Fig4J&K).

Innate cells are first responders to cutaneous wounding. Previously, it has been reported that pro-inflammatory Ly-6C^high^ macrophages accumulate at wound sites at 1 day post full-thickness wounding, contributing up to 60% of skin-resident macrophages, and gradually drop to around 10% at 7dpw^3^. Additionally, Treg depletion leads to a 6-fold increase of Ly- 6C^high^ macrophages at 7dpw, suggesting skin Tregs help transit the stages of wound healing from a pro-inflammatory to an anti-inflammatory environment. In contrast, Jag1 loss in Tregs did not lead to the same cellular skewing when compared to deletion of the entire Treg pool^3^. Instead, pro-inflammatory macrophage (defined as CD45+CD11b^high^F4/80+Ly-6C^high^Ly-6G^low^) accumulation in wounded skin remained indifferent between mice with Jag1-deficient and - sufficient Tregs, at around 6% of total macrophages at 5dpw (Figure 5A&B). Similarly, the accumulation of inflammatory Ly6C+ monocytes was unchanged (Figure 5C&D), suggesting mice with Jag1-deficient Tregs remain capable of transitioning from a pro-inflammatory to an anti-inflammatory state during wound healing. Interestingly, the absence of Jag1 in Tregs led to less neutrophil influx into wounded skin, compared to wildtype controls at 5dpw, quantified by both flow cytometry and immunofluorescence staining (Figure 5E-H). Collectively, Jag1 is unlikely to be a key factor driving the widely appreciated immunosuppressive function of skin Tregs. Rather, Jag1^pos^Tregs promote the retention of neutrophils during wound healing.

**Figure 5:**
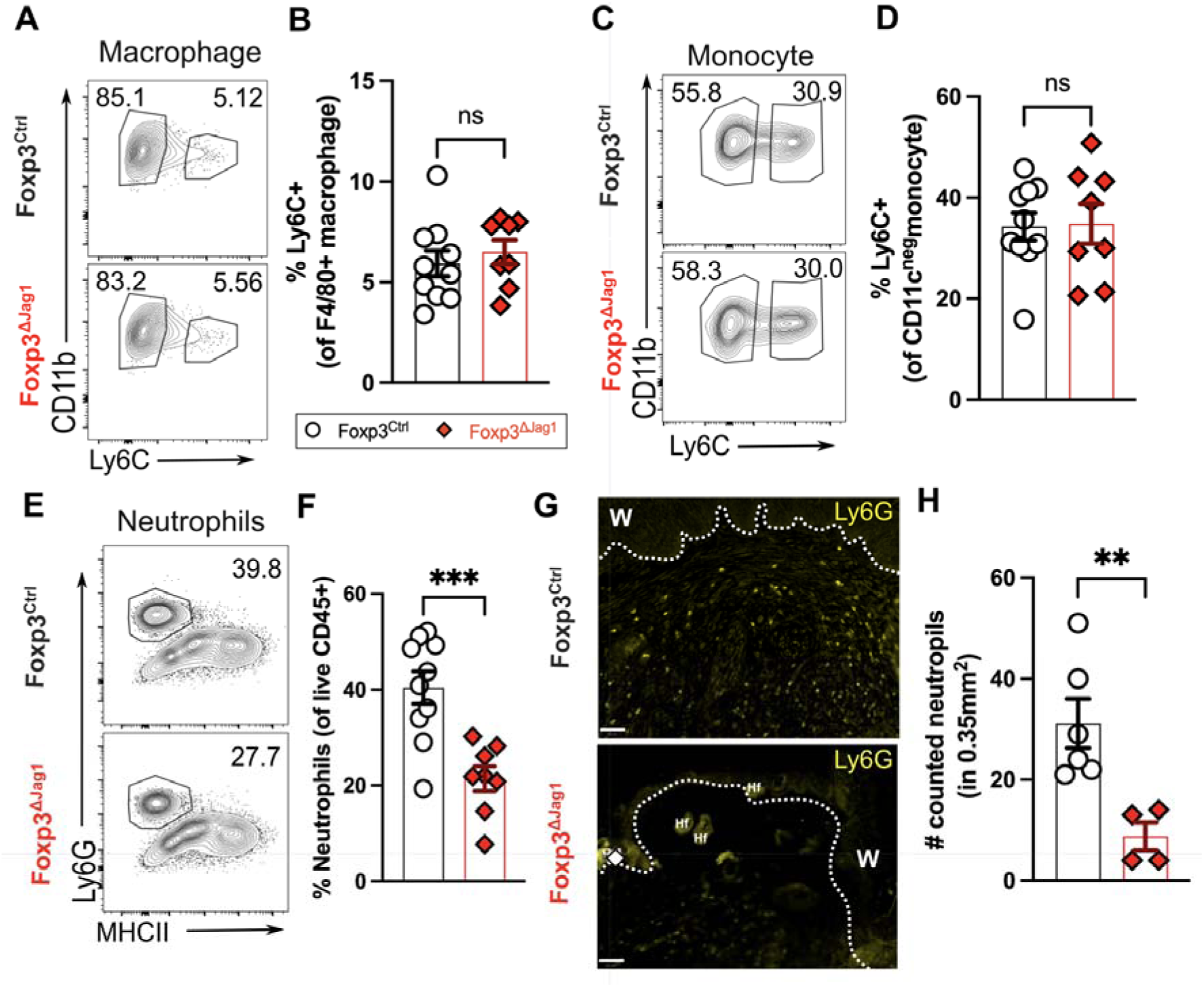
Jag1+Tregs impact neutrophil accumulation in wounded skin. Representative flow plot and quantitation of (**A** and **B**) pro-inflammatory Ly6C+ macrophages, (**C** and **D**) Ly6C+ monocytes and (**E** and **F**) neutrophils in wounded skin at 5dpw (n = 8-10 per group). (G) Representative immunofluorescence staining of Ly6G of wounded skin, with labels of wound site (w) and hair follicle (Hf). Scale bars represent 50μm. (H) Quantification of Ly6G+ neutrophils from (G) within a fixed region-of-interest. (n = 4 per group). Data were pooled from 2 independent experiments. Each individual data point represented a biological replicate, and was collectively presented with mean ± SEM. Statistics were calculated by two-way ANOVA (B, D and F) and unpaired T-test (H). ** *p <* 0.01, *** *p <* 0.001, ns = non-significant.

### Jag1+ Tregs do not Influence Re-epithelialization

Re-epithelialisation is a crucial step for successful wound healing following the inflammatory phase, by preventing excess water loss and further entry of microbial pathogens or debris. Trans-epidermal water loss measurement (TEWL) estimates moisture evaporation externally and is used quantitatively to assess epidermal integrity^28^. Besides suppressing inflammation, skin Tregs drive epidermal barrier repair by promoting bulge HFSC emigration to the epidermis, as well as their proliferation and differentiation^4^. Previous lineage tracing studies have shown that bulge cells repopulate the epidermis at 5 days post full thickness wounding^29^. These studies also indicate that bulge HFSCs are proliferative and can migrate to cutaneous wounds from 4dpw but not earlier^30^, and the derived cells can be detected in the epidermis for up to one year^31^. This indicates that while bulge HFSCs may not contribute to the immediate re-epithelisation of wounds, they play an important role in restoring the skin barrier from 4dpw onwards. Given that Jag1^pos^Tregs can drive bulge HFSC (identified as EpCam-Sca1-CD34+CD49f+) proliferation during hair regeneration^1^, we hypothesised Jag1^pos^Tregs may also regulate wound closure by mediating bulge HFSC-driven re- epithelisation.

However, mice with Jag1 deficiency in Tregs showed a similar TEWL restoration rate to baseline as wildtype controls (Supplementary Fig5A), suggesting that Jag1^pos^Tregs do not influence barrier restoration post wounding. In line with this observation, the abundance and proliferation of bulge HFSCs also remain unaffected in Foxp3^ΔJag1^ mice during wound healing (Supplementary Figure 5C&D, gating strategy in Supplementary Figure 5B), indicating that Jag1^pos^Tregs are unlikely to share the same mechanistic interaction with bulge HFSCs as observed in hair regeneration.

## DISCUSSION

Despite understanding the functional importance of skin Tregs, whether skin Tregs carry out their functions through the same molecular mechanisms as Tregs residing in other tissues is incompletely understood. One candidate is TGF-/3 signalling, recently shown to drive both hair regeneration^2^ and epithelial barrier repair ^5^. Yet, the functional importance of TGF-/3 signalling extends globally to Treg generation and/or maintenance in both lymphoid and non-lymphoid tissues (Reviewed in ^32^). Given that Jag1 is preferentially expressed in skin Tregs, we questioned whether Jag1-mediated signals are uniquely utilized by skin Tregs to drive their skin-related functions.

In the current study, we demonstrate that Jag1 is indeed preferentially expressed in skin Tregs. Jag1 expressing Tregs exhibit an activated profile, with upregulation of CD25 and CTLA4, but not ICOS, during both homeostasis and wound healing. Although it remains to be determined whether this upregulation reflects a higher immunosuppressive capacity of Jag1^pos^ Tregs, our survey of other non-Foxp3 expressing CD4+, CD8+ T cells, pro-inflammatory macrophages, or monocytes, indicates skin Tregs are unlikely to suppress *in vivo* inflammation through Jag1.

Most notably, our study has highlighted the necessity of Jag1 expression in Tregs to facilitate adequate cutaneous wound repair. Consistent with previous findings on skin Tregs, the impact of Jag1^pos^ skin Tregs is limited to a critical time window (around 5 dpw) in which wound healing transitions from the inflammatory to the proliferation phase. However, rather than suppressing inflammation, Jag1 expression in Tregs seems to drive the retention of the inflammation phase, as evidenced by neutrophil accumulation within the wounded environment. Despite their well-known tissue-damaging properties, neutrophils are crucial facilitators in wound healing. They not only dissipate infiltrating pathogens, damaged cells, and debris, but also play an increasingly appreciated role in resolving inflammation by polarizing anti-inflammatory macrophages, encouraging vascularization, and potentially promoting local cell proliferation to repair tissues (reviewed in ^33^).

Pertinently, mice lacking Notch activation in Lyz2+myeloid cells (Lyz2^cre^RBPJ^fl/fl^) show a milder inflammatory response in the heart, lung and kidney upon lipopolysaccharide exposure^34^. This is also associated with a reduction in neutrophil accumulation, reduced inflammatory cytokine detection in the liver after injury^35^, and pro-inflammatory macrophage reduction in spinal cord lesion sites after compression injury^36^. This suggests that during injury, the activation of Notch signalling in myeloid cells can promote inflammation. Mechanistically, intrinsic Notch signaling plays an important regulatory role in dampening Tregs’ own immunosuppressive functions^19^. Our study also showed that despite high CTLA4 and CD25 expression both homeostatically and during wound healing, Jag1^pos^Tregs are unlikely to self-regulate immunosuppressive capacity. While it remains to be elucidated whether Jag1^pos^Tregs directly influence Notch signalling in neutrophils, our study has uncovered an unexpected role for Jag1^pos^Tregs in modulating the inflammatory phase of cutaneous wound healing.

## Acknowledgments

We thank the Advanced Cytometry Platform (Flow Core), Research and Development Department at Guy’s and St Thomas’ NHS Foundation Trust for assistance with flow cytometry experiments.

## Author contributions

P.L.: investigation, data curation, analysis, writing – original draft, writing – Review & Editing; J.Z.X.: investigation and methodology; H.A.: investigation; M.S.: investigation; N.A. conceptualization, supervision, data curation, funding acquisition, writing – original draft, writing – Review & Editing.

## CONTACT FOR REAGENT AND RESOURCE SHARING

Requests for reagents and resources should be directed to the Lead Contact, Niwa Ali

## Methods

### Animal study design

Wildtype C57BL/6 and FoxP3^eGFP-CreERT2^ mice (JAX: 016961) were purchased from Charles River, and Jag1^fl/fl^ (JAX: 010618) were gift from Prof Rosenblum in UCSF. FoxP3^eGFP-CreERT2^ were crossed to Jag1^fl/fl^ to generate FoxP3^eGFP-CreERT2^ Jag1^fl/fl^ in King’s College London BSU NHH animal unit. Mice were maintained through routine breeding, were fed with a standard chow diet and housed in line with UK regulations. All experiments were performed on animals with no prior procedures, with authorisation under the project license PP6051479. Littermates of the same sex were genotyped and assigned to experimental groups based on genotyping results.

Tamoxifen (Sigma, T5648-5G) were sonicated in 37^°^C water bath and dissolved in corn oil, at 2.5mg/ml concentration. For conditional Jag1 knockout, 7-10 week old Foxp3^eGFP-Cre-ERT2^Jag1fl/fl and Foxp3^eGFP-Cre-ERT2^Jag1fl/wt or wt/wt were injected with tamoxifen intraperitonially at 75-100mg/kg in the indicated interval before harvesting. All characterisation and steady state experiments were performed on mice of mixed gender, and only females were used for further characterisation in wound healing experiments.

### Skin wounding assays and analysis

Mice were anaesthetised by inhalation of vaporized 1.5% isoflurane, shaved and subcutaneously injected with Vetersgesics. Two full thickness excisional wounds were made on dorsal back of mice under and anaesthesia, using a 4-mm biopsy punch (Stifel Laboratory Research). Wounds were photographed daily till harvest. The same ruler was placed next to wound area for measurement standardization. Wound area was measured using ImageJ^37^(Fiji NIH), and closure ratio of each time point was calculated relative to wound area at 0dpw.

Transepidermal water loss (TEWL) of each wound was measured with Tewameter TM 300 probe (Courage + Khazaka electronic GmbH) according to the manufacturer’s protocols. Measurement were made on 0dpw (immediately after excision wound) and every 24 hours thereafter. Each datapoint is an average of four TEWL measurements.

### Tissue Processing

Whole murine dorsal skin was finely minced with scissors and digested in 500ul digestion medium per cm^2^ of skin. Digestion medium was prepared with 2mg/ml collagenase (Sigma), 0.1mg/ml DNase (Sigma) and 0.5mg/ml hyaluronidase (Sigma), dissolved in C10 medium [10% FBS, 1% Pen/Strep, 1 mM Na-pyruvate, 1% HEPES, 1% non-essential amino acid, 0.5% 2-mercaptoethanol in RPMI-1640 with L-glutamine medium]. After 45 min incubation at 37^°^C 255rpm, single cell suspension was washed with 20ml C10 medium, and filtered through 100um then 40um cell strainer. Lymph nodes were mechanically smashed, washed with FACS buffer [2% Fetal calf serum, 1mM EDTA and 0.1% sodium azaide in PBS], and filtered through 70um cell strainer. Epidermal cells were prepared by floating skin on 0.5%Trypsin-EDTA (Thermofisher) for 1hr at 37^°^C, before gently removed from dermal part and washed with C10 medium. Lung was finely minced and digested with 1mg/ml Collagenase A (Sigma) and 0.1mg/ml DNase I (Sigma) in R10 [10% FBS and 1% Pen/Strep in RPMI-1640 medium]. After 1hr at 37^°^C 180rpm, single cell suspension is passed through 70um filter, spin for 5 min at 4^°^C 1800rpm, and treated with 500ul ACK Lysing Buffer for 30s to 1min before washed with 1xPBS. All single cell suspensions were then centrifuged at 1800rpm at 4^°^C for 4 min, and resuspended in 1m FACS buffer. Total live cells were determined using NucleoCounter NC-200 (Chemometec) in 1:20 dilution, before downstream process.

For histology, skin tissue was fixed in 10% formalin overnight, followed by PBS washes, stored at 70% Ethanol overnight, and embedded in paraffin. Each sample was cut and levelled to wound area, before manually assembled as tissue microarray blocks using a 7- mm biopsy punch. 5um sections were cut, mounted and sent to Tissueplexia (Scotland) for multiplex-staining using anti-FoxP3 (14-5773-82, Thermofisher) and anti-Ly6G (127602, Biolegend). H&E were performed according to manufacturer’s instruction and imaged using a Nanozoomer (Hamamatsu photonics) with a ×40 objective.

### Flow cytometry

For flow cytometry staining, 1.5-4 million cells per condition were plated in round bottom 96 well plate, and stained with 50ul of stated surface antibodies on ice for 20 min. After washed with FACS buffer, cells were fixed and permeabilised by FoxP3/Transcription Factor Staining buffer set (eBioscience) on ice for another 20 min, before washed with permeabilization buffer and lastly stained with intracellular antibodies (resource table), again on ice for 20 min. Samples were run on Fortessa LSRII (BD Bioscience) in KCL BRC Flow Cytometry Core. For compensation, UltraComp eBeads^TM^ (Thermofisher) were stained with each surface and intracellular antibody following the same cell staining protocol. ArC^TM^ Amine Reactive Compensation Bead Kit (Thermofisher, A10346) were used for GhostDye^TM^ Live/Dead stain. All gating and data analysis were performed using FlowJo v10, while statistics were calculated using Graphpad Prism 10.

### RNA isolation and quantification PCR

Lymph node single cell suspension was resuspended in pre-sort medium [2% FBS, 1% Pen/Strep, 2 mM EDTA, 25mM HEPES in RPMI-1640 without phenol red]. CD45+ CD3+CD4+CD25high (Tregs), CD45+CD3+CD4+CD25neg(Teffs) and CD45+CD3+CD8+ cells were sorted using FACSAria™ Fusion Flow Cytometer (BD) with 100um nozzle in KCL BRC Flow Cytometry Core. Cells were sorted into RPMI-1640 supplemented with 10% heat-inactivated FBS and 1% Pen/Strep, and spun at 300g for 10min at 4^°^C. After removal of supernatant, cell pellet were snap-frozen in liquid nitrogen and stored at -80^°^C. RNA was extracted using NucloSpin RNA XS, Micro kit for RNA purification (740902, Macherey-Nagel) according manufacturer’s instruction. RNA integrity and concentration were then determined by RNA 6000 Pico Kit on Bioanalyzer (Agilent). RNA were then normalized and synthesied into cDNA using iScript cDNA synthesis kit (Bio-Rad). Quantitative PCR were performed using TaqMan^TM^ PreAmp Master Mix Kit (ThermoFisher) on 384-well plate according to manufacturer’s instruction, run with 4 technical replica each condition. The following TaqMan probes were used: Gapdh (Mm99999915_g1, VIC), Foxp3 (Mm00475162_m1, FAM) and Jag1 (Mm00496904_m1, FAM), Plates were then run on CFX384 Touch Real-Time PCR system (BioRad).

## QUANTIFICATION AND STATISTICAL ANALYSIS

Parameters such as sample size, dispersion or precision are reported in Figure Legends. Statistical analyses were performed in Prism 10.1 (GraphPad). Details of the statistics and appropriate test used are also indicated in Figure Legends. *p < 0.05, **p < 0.01 ***p < 0.001, ****p < 0.001, p-values greater than 0.05 was identified as not statistically significant.

### Bulk RNAseq reanalysis

Read-count tables were obtained from NCBI database: GSE182322 (Ref ^10^), and downstream processed according to presented method. Each data point in Figure 1 represents the sum of both gene raw counts from ST2- and ST2+ Tregs per mouse.

| REAGENT or RESOURCE | SOURCE | IDENTIFIER |
| --- | --- | --- |
| **Antibodies** |  |  |
| Mouse T Cell Panel |  |  |
| CD45 Monoclonal Antibody (30-F11), Alexa Fluor 700, eBioscience™ | ThermoFisher | Cat# 56-0451-82, RRID: AB_891454 |
| CD3 Antibody, anti-mouse, PerCP-Vio® 700, REAfinity™ | Miltenyi Biotec | Cat# 130-120-826, RRID: AB_2752207 |
| TCRγ/δ Antibody, anti-mouse, PE-Vio® 770, REAfinity™ | Miltenyi Biotec | Cat# 130-123-290, RRID: AB_2802022 |
| Brilliant Violet 650™ anti-mouse CD4 Antibody | BioLegend | Cat# 100545, RRID: AB_11126142 |
| CD8a Antibody, anti-mouse, APC-Vio® 770, REAfinity™ | Miltenyi Biotec | Cat# 130-120-806, RRID: AB_2752203 |
| CD25 Antibody, anti-mouse, PE-Vio® 615, REAfinity™ | Miltenyi Biotec | Cat# 130-123-028, RRID: AB_2811434 |
| BD Horizon™ BV605 Hamster Anti-Mouse CD27 | BD Biosciences | Cat# 563365, RRID: AB_2738160 |
| Jagged1 Antibody (E-12) | Santa Cruz Biotechnology | Cat# Sc-390177, RRID: AB_2892141 |
| Anti-Mouse CD339 (Jagged 1) PE | eBioscience | Cat# 12-3391-80, RRID: AB_1963616 |
| Brilliant Violet 711™ anti-human/mouse/rat CD278 (ICOS) Antibody | BioLegend | Cat# 313547, RRID: AB_2734288 |
| BD Horizon™ BV786 Mouse Anti-Ki-67 | BD Biosciences | Cat# 563756, RRID: AB_2732007 |
| FoxP3 Antibody, anti-mouse, Vio® R667, REAfinity™ | Miltenyi Biotec | Cat# 130-111-682, RRID: AB_2651771 |
| Brilliant Violet 421™ anti-mouse CD152 Antibody | BioLegend | Cat# 106311, RRID: AB_10901170 |
| Anti-Mouse/Rat Foxp3 eFluor 450 | eBioscience | Cat# 48-5773-82 |
| Anti-Mouse CD152 (CTLA4) PE | BD | Cat# 553720 |
| Anti-Human Ki67 PE-Cy7 | BD | Cat# 561283 |
| Anti-Mouse TCR gd<br>PerCP-Cy 5.5 | Biolegend | Cat# 118117 |
| Anti-Mouse CD45<br>Alexa Fluor 700 | eBioscience | Cat# 56-0451-82 |
| Anti-Mouse CD25<br>APC-eFluor 780 | eBioscience | Cat# 47-0251-82 |
| Brilliant Violet 605<br>anti-mouse CD8a<br>Antibody | Biolegend | Cat# 100743 |
| Brilliant Violet 650™<br>anti-mouse CD4<br>Antibody | Biolegend | Cat# 100545 |
| Brilliant Violet 711<br>anti-mouse CD3<br>Antibody | Biolegend | Cat# 100241 |
| Mouse Myeloid Panel |  |  |
| BD Horizon™<br>BUV395 Rat Anti-Mouse Ly-6G | BD<br>Biosciences | Cat# 565964, RRID: AB_2716852 |
| Brilliant Violet 421™<br>anti-mouse I-A/I-E<br>Antibody | BioLegend | Cat# 107632, RRID: AB_2650896 |
| Brilliant Violet 510™<br>anti-mouse/human<br>CD45R/B220<br>Antibody | BioLegend | Cat# 103248, RRID: AB_2650679 |
| Brilliant Violet 510™<br>anti-mouse CD3<br>Antibody | BioLegend | Cat# 100234, RRID: AB_2562555 |
| Brilliant Violet 650™<br>anti-mouse CD11c<br>Antibody | BioLegend | Cat# 117339, RRID: AB_2562414 |
| Brilliant Violet 785™<br>anti-mouse F4/80<br>Antibody | BioLegend | Cat# 123141, RRID: AB_2563667 |
| FITC anti-mouse Ly-6C Antibody | BioLegend | Cat# 128005, RRID: AB_1186134 |
| CD207 (Langerin)<br>Monoclonal Antibody<br>(eBioL31), PE,<br>eBioscience™ | ThermoFisher | Cat# 12-2075-82, RRID: AB_763452 |
| CD45 Monoclonal<br>Antibody (30-F11),<br>Alexa Fluor 700,<br>eBioscience™ | ThermoFisher | Cat# 56-0451-82, RRID: AB_891454 |
| APC/Cyanine7 anti-mouse/human CD11b<br>Antibody | BioLegend | Cat# 101225, RRID: AB_830641 |
| **Epidermal panel** |  |  |
| BD Horizon™<br>BUV395 Rat Anti-Mouse CD45 | BD<br>Biosciences | Cat# 564279 |
| CD326 (EpCAM)<br>Antibody, anti-mouse,<br>REAfinity™ | Miltenyi | Cat# 130-117-871 |
| BD Horizon™ BV786<br>Mouse Anti-Ki-67 | BD<br>Biosciences | Cat# 563756 |
| BD Horizon™ BV650<br>Rat Anti-Human<br>CD49f | BD<br>Biosciences | Cat# 563707 |
| Alexa Fluor® 700<br>anti-mouse I-A/I-E<br>Antibody | Biolegend | Cat# 107622 |
| CD207 (Langerin)<br>Antibody, anti-mouse,<br>PE-Vio® 615,<br>REAfinity™ | Miltenyi | Cat# 130-112-994 |
| <u>BD Pharmingen™</u><br><u>Alexa Fluor® 647 Rat</u><br><u>anti-Mouse CD34</u> | BD<br>Biosciences | Cat# 560230 |
| Sca-1 Antibody, anti-mouse, REAfinity™ | Miltenyi | Cat# 130-106-220 |
| **Chemicals, peptides, and recombinant proteins** |  |  |
| **Tamoxifen** | Sigma-Aldrich | Cat# T5648 |
| Collagenase | Sigma-Aldrich | Cat #C9407 |
| DNase | Sigma-Aldrich | Cat #DN25 |
| Hyaluronidase | Sigma-Aldrich | Cat #H3506 |
| Collagenase A | Sigma-Aldrich | Cat# **10103578001** |
| DNase 1 | Sigma-Aldrich | **Cat# 10104159001** |
| ACK Lysing Buffer | ThermoFisher | **Cat#** A1049201 |
| Fetal Bovine Serum | ThermoFisher | Cat# 10500064 |
| Penicillin-Streptomycin (5,000<br>U/mL) | ThermoFisher | Cat# 15070063 |
| Sodium Pyruvate<br>(100 mM) | ThermoFisher | Cat# 11360070 |
| HEPES (1M) | ThermoFisher | Cat# 15630056 |
| MEM Non-Essential<br>Amino Acids Solution<br>(100X) | ThermoFisher | Cat# 11140050 |
| 2-mercaptoethanol | ThermoFisher | Cat# 21985023 |
| RPMI-1640 medium | ThermoFisher | Cat# 11875093 |
| UltraPure™ 0.5M<br>EDTA, pH 8.0 | ThermoFisher | Cat# 15575020 |
| PBS, pH 7.4 | ThermoFisher | Cat# 10010023 |
| FoxP3/Transcription Factor Staining buffer set | eBioscience | Cat# 00-5523-00 |
| UltraComp eBeads | Thermofisher | Cat# 01-2222-42 |
| ArCTM Amine Reactive Compensation Bead Kit | Thermofisher | Cat# A10346 |
| Zombie UV™ Fixable Viability Kit | BioLegend | Cat# 423107 |
| **Deposited data** |  |  |
| Bulk RNAseq data | Spath et al(13) | GEO: GSE182322 |
| **Experimental Models: Organisms/Strains** |  |  |
| WT C57BL/6J | Charles River | RRID: IMSR_CRL:027 |
| Foxp3<sup>eGFP-Cre-ERT2</sup><br>(*Foxp3*<sup>tm9(EGFP/cre/ERT2)</sup><br>*Ayr/J*) C57BL/6 | The Jackson Laboratory | RRID: IMSR_JAX:016961 |
| Jag1<sup>fl/fl</sup> (B6.129S-<br>*Jag1*<sup>tm2Grid/J</sup>) | The Jackson Laboratory | RRID: IMSR_JAX:010618 |
| **Software and Algorithms** |  |  |
| FlowJo (version 10.8.1) | Becton Dickinson | https://www.flowjo.com/solutions/flowjo;<br>RRID:SCR_008520 |
| Fuji (Image J) | Fuji (Image J) | https://fiji.sc |
| NDP.view2 Image viewing software (U12388-01) | Hamamatsu | https://www.hamamatsu.com/us/en/product/life-science-and-medical-systems/digital-slide-scanner/U12388-01.html<br>RRID:SCR_025177 |
| QuPath (version 0.5.0-arm64) | Bankhead et al. | https://qupath.github.io/ |
| Prism (version 10.1.0) | GraphPad Software | http://www.graphpad.com/;<br>RRID:SCR_002798 |
| Inkscape (version 1.3.2) | Inkscape | https://inkscape.org/;<br>RRID:SCR_014479 |
| R (version 4.3.2) | CRAN | https://www.r-project.org/ |
| RStudio (version 2021.09.0) | RStudio | https://www.rstudio.com/ |
| R package: edgeR<br>(version 4.0.16) | Robinson et al<br>(38) | http://bioconductor.org/packages/edgeR/ |
|  |  | RRID:SCR_012802 |
| R package: limma<br>(version 3.58.1) | Ritchie et al | http://bioinf.wehi.edu.au/limma/ |
|  |  | RRID:SCR_010943 |
| R package:<br>org.Mm.eg.db | Carlson M. | https://bioconductor.org/packages/org.Mm.eg.db/ |
| (version 3.58.1) |  | RRID:SCR_023488 |
| Other |  |  |
| NanoZoomer |  |  |
| BD FACSAria II Cell<br>Sorter |  | --- |
| BD LSR-Fortessa<br>analyzer |  |  |
| 4-mm biopsy punch | Stifel |  |

## SUPPLEMENTARY FIGURES

**Figure S1.**
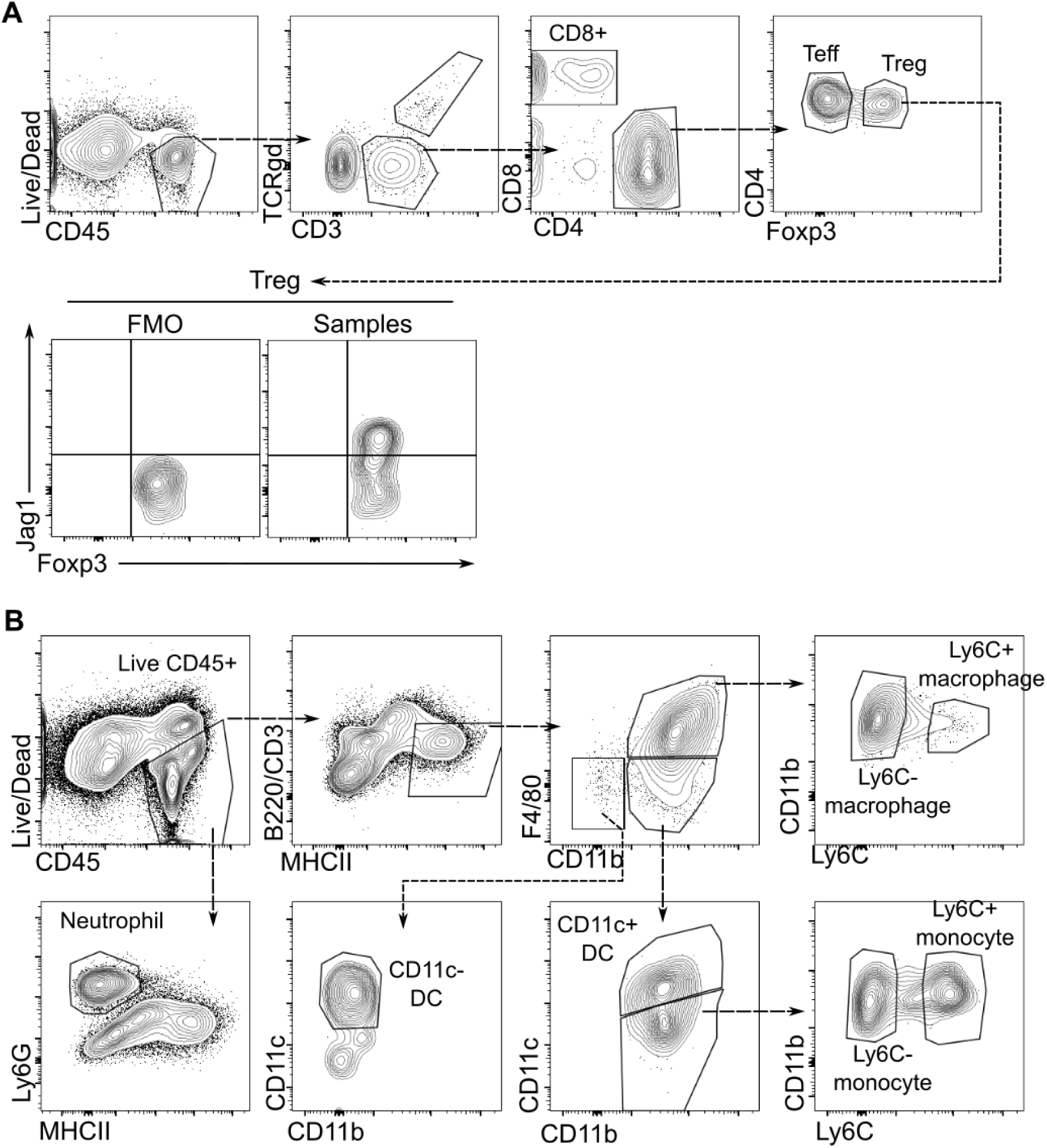
Flow cytometric gating strategy to identify Jag1+Tregs and skin T cell populations. Cells were first stained with Zombie UV live/dead stain and CD45 to identify live CD45. **(A)** T cell panel. T cells (TCRgd-CD3+) populations were further segregated based on CD8 and CD4 expression. Tregs were gated based on their Foxp3 expression. Jag1 gate was further determined based on FMO of all samples within each experiment. **(B)** Myeloid panel. Neutrophils (Ly6G+MHCII-) were gated directly from live CD45+ gate. B-cells and T-cells were excluded using B220 and CD3. Within the B220-CD3-MHCII+ gate, macrophages were identified as F4/80+ CD11b+ population, and were further separated based on their Ly6C expression. Dendritic cells were separated into CD11b+CD11c+ and CD11b-CD11c+ populations. The remaining CD11b+CD11c-population was classified as monocytes, which was further subset into Ly6C+ and Ly6C-populations.

**Figure S2.**
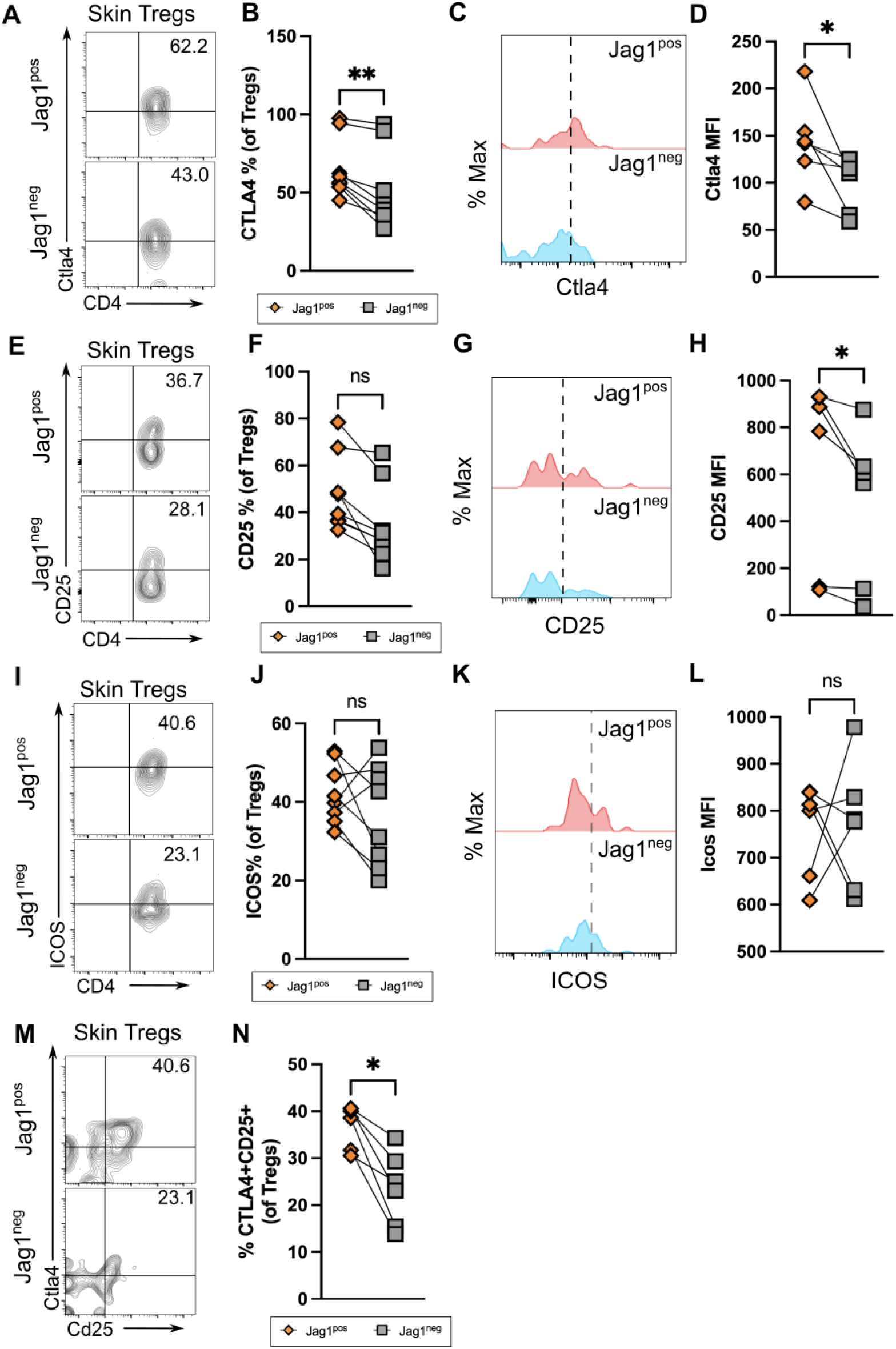
Jag1+Tregs are phenotypically more activated than Jag1-Tregs. Representative flow plot and quantitation of % Ctla4 (**A** & **B**), %CD25 (**E** & **F**), %ICOS (**I** & **J**) and % Ctla4+Cd25+ (**I** and **J**) in Jag1^pos^ and Jag1^neg^ skin Tregs from wildtype mice. Representative histogram and MFI quantification of Ctla4 (**C** & **D**), CD25 (**G** & **H**) expressed in Jag1^pos^ and Jag1^neg^ skin Tregs. Dash line of histogram indicates gating from negative controls. Data were pooled from 2 independent experiments with n = 6. Each paired data point represented a biological replicate. Statistics were calculated by paired Wilcoxon t-test. * p < 0.05, ** p <0.01.

**Figure S3.**
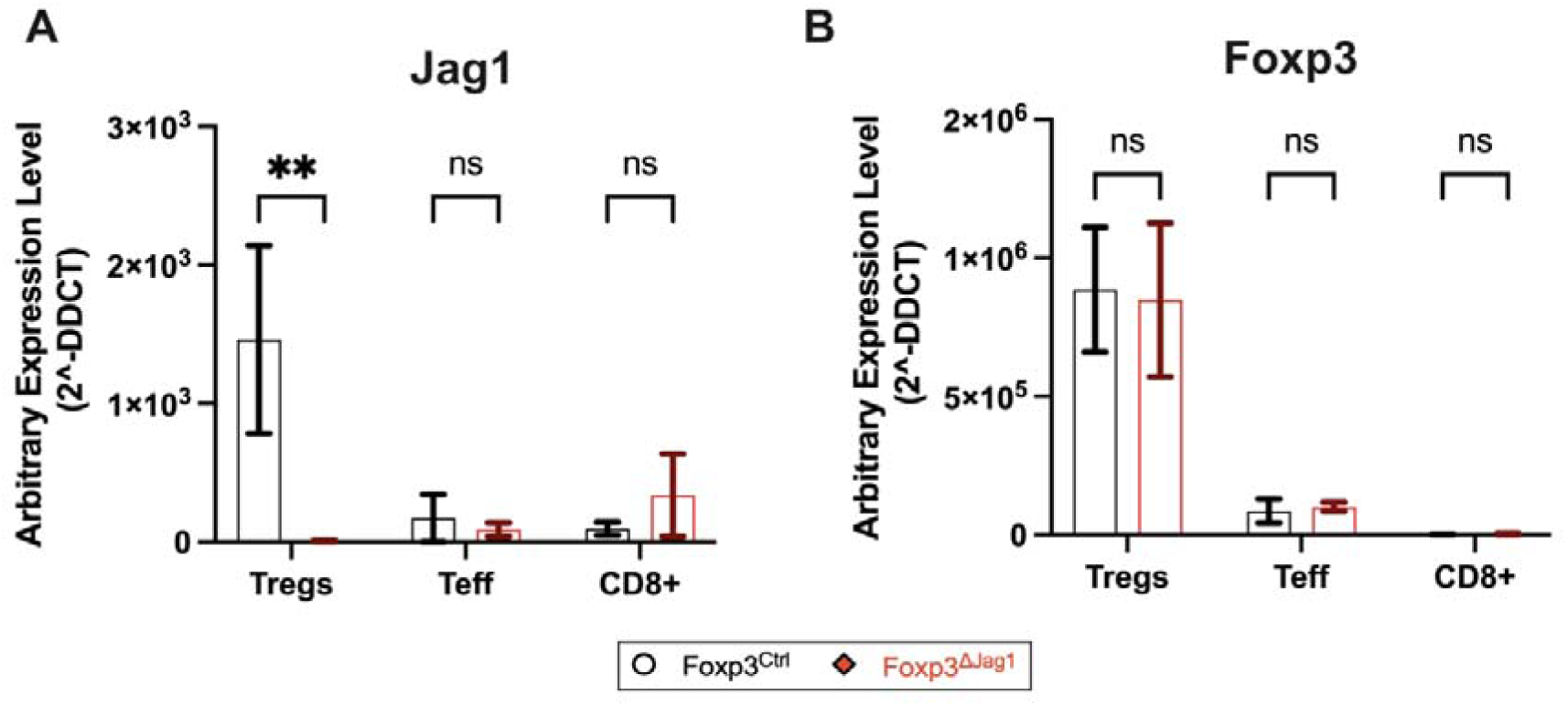
qPCR of sorted Tregs to confirm Jag1 deletion in Tregs. Data were pooled from 2 independent experiments with n = 3-4. Results were presented as mean ± SEM. Statistics were calculated by two-way ANOVA, ** *p <* 0.01, ns = non-significant.

**Figure S4:**
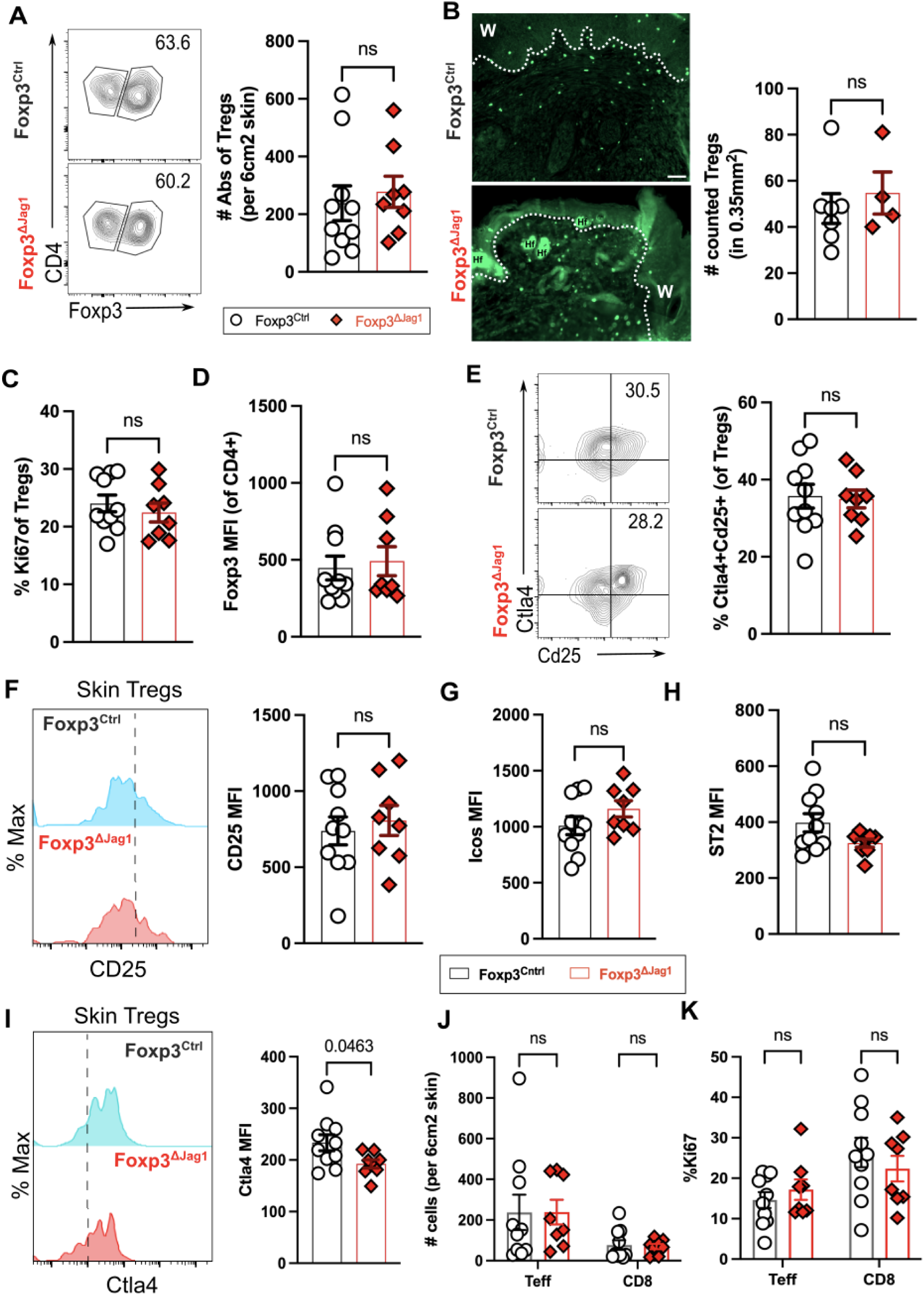
Jag1+Tregs do not influence T cell accumulation in wounded skin. (**A**)Representative flow plots and absolute abundance of Tregs from wounded skin at 5dpw (n = 8-10 per group). (B) Representative immunofluorescence staining and quantification of Foxp3 in wounded skin, with labels of wound site (w) and hair follicle (Hf). Scale bars represent 50μm. (n = 4 per group). Flow cytometric quantification of (C) Treg %Ki67 and (D) mean fluorescence intensity (MFI) of Foxp3 in CD4+ T cells (n = 8-10 per group). Representative flow plot or histogram and quantification of (E) %Ctla4+CD25+, MFI of (F) CD25, (G) Icos, (H) ST2 and (I) Ctla4 in Tregs of wounded skin (n = 8-10 per group). Quantification of **(J)** Tregs, Teffs and CD8+ T cells abundance, and **(K)** % Ki67 of indicated T cells residing in wounded skin. Data were pooled from 2 independent experiments. Each individual data point represented a biological replicate, and was collectively presented with mean ± SEM. Statistics were calculated by unpaired T-test (A-I) and two-way ANOVA (J and K). ns = non-significant.

**Figure S5:**
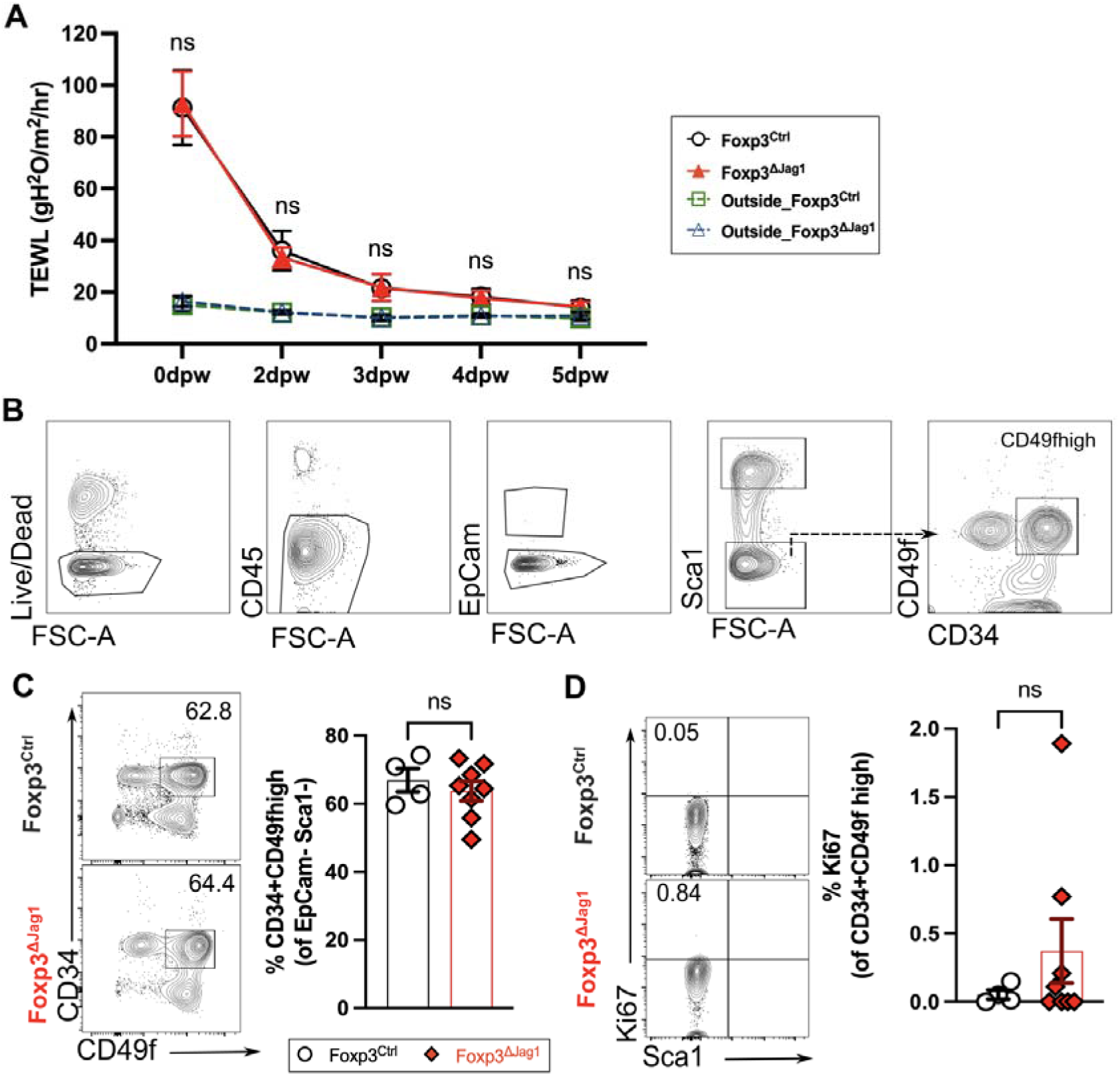
Jag1+Tregs do not influence HFSC homeostasis during wound healing. **(A)** Kinetics of trans epidermal water loss (TEWL) taken during wound healing of Foxp3^Ctrl^ and Foxp3^ΔJag1^, with skin outside wound site setting the baseline of intact skin barrier (n = 8 per group).**(B)** Flow cytometric gating strategy to identify epithelial cell populations in skin. Cells were stained with Zombie UV live/dead stain and CD45 to identify live CD45neg populations. Epithelial cells were further separated into bulge HFSCs (Sca1+CD49f+CD34+)). Representative flow plots and quantification of bulge HFSCs abundance (C) and their proliferation (D)from wounded skin at 5dpw (n = 4-8 per group). Data were pooled from 2 independent experiments. Each individual data point represented a biological replicate, and was collectively presented with mean ± SEM Statistics were calculated by two-way ANOVA (A) and unpaired T-test (C and D). ns = non-significant.. Statistics were calculated by unpaired T-test (B), and two-way ANOVA (B, D and F). ns = non-significant.

